# Layers to Leaves: A Suite of Modular 3D Printed Hydroponics Components for Research and Education

**DOI:** 10.1101/2025.07.24.666580

**Authors:** E. Shaw, S.K. Chandramouli, M.P. Dzakovich

**Affiliations:** USDA-ARS Children’s Nutrition Research Center, Department of Pediatrics, Baylor College of Medicine, Houston, TX, 77030, USA; University of Houston Department of Biomedical Engineering, Houston, TX, 77204, USA; Rice University Department of Biosciences, Houston, TX, 77251, USA

**Keywords:** Plant Biology, STEM, Horticulture, 3D Printing, Salinity, Spinach

## Abstract

Hydroponics is a widely utilized technique to precisely control the plant growing environment and maximize productivity. In some cases, hydroponics systems can be expensive and require specialized expertise to build and maintain. Here, we present a suite of 3D-printed devices that can be constructed into a single or double tower hydroponics system. This system is easily scalable to fit the needs of a user and can be implemented in a variety of research and educational contexts. Components in this suite can be made from inexpensive plastic filament using household-grade 3D printers. The vertical design of these systems allows for users to maximize space-use-efficiency in growth chambers, greenhouses, or even classrooms. We describe the construction of our system and provide example data from a validation study growing spinach (*Spinacia oleracea L.*) under different salinity conditions. This suite of 3D printed components can be utilized by researchers and educators alike to capitalize on the benefits of hydroponics in a flexible, budget-friendly way.

## Introduction

3D printing is an emerging form of additive manufacturing that allows for the creation of custom parts tailored to meet the needs of an end user. Although metal-based 3D printers are utilized in high-end industrial manufacturing, most 3D printers available to consumers use inexpensive plastic filament (e.g. polylactic acid; PLA) that is widely available online or in retail stores [1]. Manufacturing advances have allowed the 3D printer industry to generate a range of product lines available at a low cost to consumers; many entry level devices cost around $300. As such, this technology is quickly being adopted in research settings [1–4] as well as incorporated into classroom curriculum to promote STEM education [5–8]. One application of 3D printing is manufacturing modular hydroponic systems that can be adjusted to the needs of the end user without the need for complete refabrication.

Hydroponics is a widely used approach to maximize plant growth, development, and uniformity. Plants grown hydroponically benefit from optimized mineral nutrition, absence of water stress common in field-grown crops, and are often grown in systems that generally control other environmental factors (e.g. vertical farms) [9,10]. While hydroponics has long been adopted by industry to maximize plant yield, it continues to have innumerable applications for research. Mineral nutrition studies can be easily conducted using hydroponics due the customizable nature of nutrient solutions and minimized nutrient adsorption/desorption that commonly occurs with solid substrates and soil [11]. By making mineral nutrition consistent among plants, the effects of plant genotype as well as environmental variables can be studied in relative isolation compared to soil or field-grown plants [12–14]. Hydroponics also accommodates intrinsic labeling using precious radiolabeled or nonradioactive isotopes that would be lost to a solid substrate [11]. These experiments provide deeper insights into nutrient turnover and partitioning as well as their bioavailability in nutritional studies. While there are countless uses for hydroponics in industry and research, this technique is increasingly being utilized as a means of STEM education in school curriculum.

Hydroponics can be utilized in educational settings for a wide range of student ages to provide hands-on learning experiences in plant biology [15–18]. Limited studies indicate that gardening and hydroponics can improve children’s attitudes towards fruit and vegetable consumption [19–22] as well as their perceptions of science, technology, engineering, and math (STEM) disciplines [16,17,23,24]. Similarly, 3D printing is also quickly becoming integrated into school curriculum for elementary school to university aged individuals [5–8]. There is enormous potential to leverage 3D printing to create modular, user-friendly hydroponics systems for both research and educational purposes.

We sought to develop a suite of 3D printed objects that can be assembled into a cost-effective and modular hydroponics system. Components from this suite of models can also be used independently of the systems we designed (e.g. plant cups). While the components are meant to work in concert with one another, these objects can serve as scaffolds for in any number of downstream applications. Here, we describe the components of the system, its construction, and expected outcomes using spinach (*Spinacia oleracea L.*) as a proof of concept.

## Materials and Methods

### System Description

The modular hydroponics system consists of three main parts: 1) the nutrient solution bin; 2) one or more 3D printed tower(s); 3) the 3D printed lid. The system is designed to support either a Single or Double Tower configuration; maximizing flexibility for various end users (e.g., researchers, educators) while remaining cost-effective using 3D printing. Components are easily interchangeable (Tables 1-2), allowing users to adapt or integrate parts into other hydroponic setups as needed. Files for 3D components can be found in the NIH3D database for the Single (https://3d.nih.gov/entries/3DPX-021941) and Double (https://3d.nih.gov/entries/3DPX-021942) tower systems [25,26].

**Table 1.** Tower modules for the single tower (ST) system.

| Part Number | File Name | Part Name | Count for one system |
| --- | --- | --- | --- |
| 1 | 4-Cup_Planting_Module.stl | 4-cup ST Planting Module | Variable |
| 2 | Tower_Lid.stl | Tower Lid | 1 |
| 3 | Tower_Distributor.stl | Tower Distributor | 1 |
| 4 | Stream_Breaker.stl | Stream Breaker | 1 |
| 5 | Tower_Short_Spacer.stl | Tower Short Spacer | Variable |
| 6 | Standard_Pot.stl | Standard Pot | 4 x 4-cup ST Planting Module |
| 7 | Reservoir_Tower_Adaptor.stl | Tower Adaptor | 1 |

**Table 2.** Lid modules for the single tower (ST) system.

| Part Number | File Name | Part Name | Count for one system |
| --- | --- | --- | --- |
| 8 | Reservoir_Spacer_Long_Edge.stl | Reservoir Spacer Long Edge | 2 |
| 9 | Reservoir_Lid_Left.stl | Reservoir Lid Left | 2 |
| 10 | Reservoir_Lid_Right_Access_Hole.stl | Reservoir Lid Right Access Hole | 1 |
| 11 | Reservoir_Lid_Right.stl | Reservoir Lid Right | 1 |
| 12 | Reservoir_Spacer_Short_Edge.stl | Reservoir Spacer Short Edge | 2 |
| 13 | Access_Hole_Cap | Access Hole Cap | 1 |
| 14 | Not applicable | M5 Screw and nut - 25mm | 6 |

### Nutrient Solution Bin and Pump

The hydroponics system was designed to work with ULine S-20588GR 20L Round Trip Totes (ULine; Pleasant Prairie, Wisconsin, United States). These bins, measuring 21.8" × 15.2" × 9.3", are made of high-density polyethylene, a durable, chemically inert material, and are rated to support up to 70 lbs. To inhibit algal growth, a dark-colored bin (e.g., gray) was used to minimize light penetration into the nutrient solution.

Each bin also houses an ActiveAqua Submersible Pump 250 (Hydrofarm; Petaluma, California, USA) located in the center, which circulates nutrient solution from the reservoir to the top of the planting towers. Similar pumps, or even external pumps, may be utilized by the user at their discretion. Structural modifications to the system would need to be made to accommodate an external pump.

### 3D Printed Towers

To improve growing capacity beyond what a horizontal growth-system allows, our system capitalizes on vertically stacked planting towers. Each tower is built using modular 4-cup units, available in 3- and 5-cup variants, which are stacked with spacer modules in either standard or extended lengths (Figure 1; Figure 2, Detail B). These variable planting modules have a finite number of spaces for plant cups (Figure 2, Detail C) that allow for 2” neoprene cloning collars to support plants throughout their lifecycle. Single-tower plant modules, cups, spacers, and stream breaker, are remixed from Thingiverse user ‘boundarycondition’ [27].

**Figure 1.**
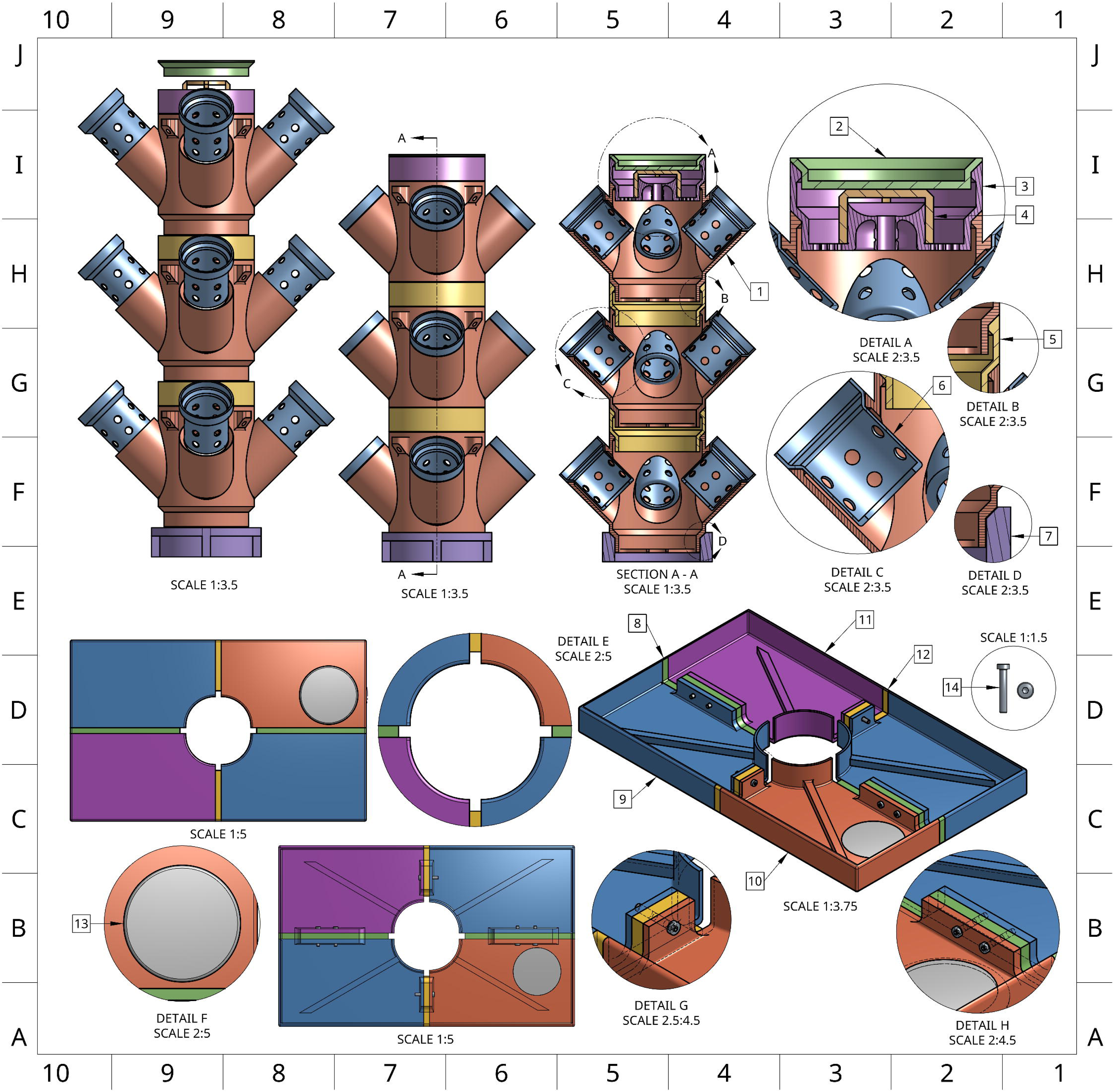
Multi-perspective rendering of the Single Tower system highlighting lid and tower components. An interactive 3D rendering of the Single Tower system (https://sketchfab.com/3d-models/one-tower-enviroment-696d529bdf3249bb8fe916f2aff49538) is available for close examination of parts in the context of a completed system.

**Figure 2.**
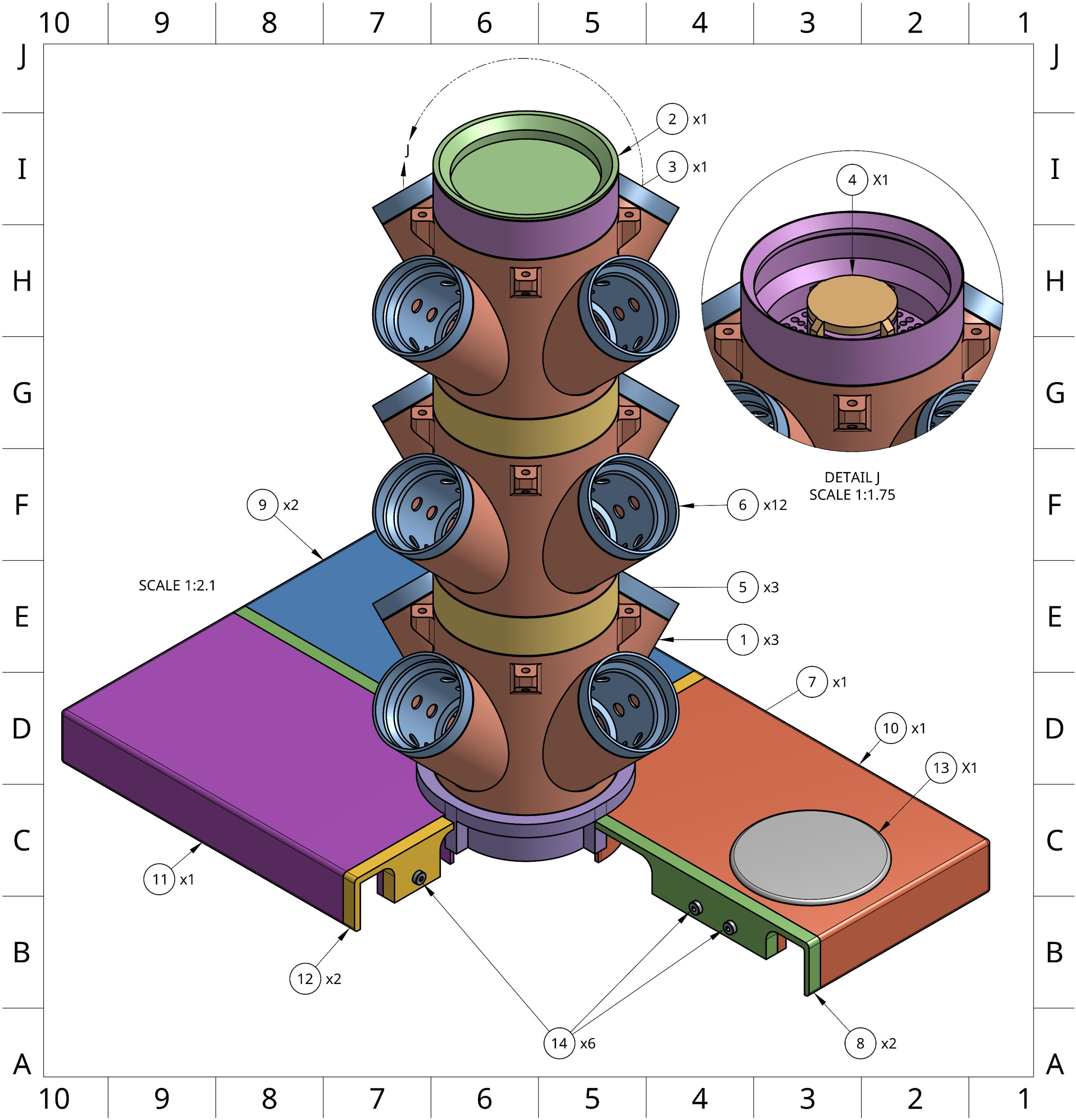
Detailed rendering of the Single Tower system showing required components necessary for assembly with the exception of one lid piece for a clearer view of the tower-lid interface. An interactive 3D rendering of the Single Tower system (https://sketchfab.com/3d-models/one-tower-enviroment-696d529bdf3249bb8fe916f2aff49538) is available for close examination of parts in the context of a completed system.

All modules are designed with a central channel (Figure 2, Section A-A) for routing ½” internal-diameter flexible vinyl tubing, which runs up to a final distributor at the top of the tower. This tubing is secured by the distributor’s nipple connector, and a stream breaker is installed over the distributor opening to diffuse flow (Figure 2, Detail A). The distributor is then capped and recommended to be held in place with a small (∼500g) weight on the lid or sealed with silicone. Once transported to the top of the tower, the nutrient solution exits through a radial array of drainage holes, evenly cascading down through the tower, irrigating the roots, and returning to the reservoir. This circulation also promotes oxygenation of the root zone to encourage healthy plant growth.

### 3D Printed Lid

To interface the towers with the ULine totes, a custom multi-part lid was designed, consisting of modular lid panels, spacer components, and an access cap (Figure 2). These parts are secured using M5 screws and nuts (Figure 2, Detail G/H). This bolting configuration, in conjunction with the radially symmetrical tower adaptor (Figure 2, Detail E), ensures mechanical stability under the compressive force of a loaded tower (Figure 2). To minimize potential leaks and reduce the risk of microbial contamination, all seams are sealed with silicone and optionally wrapped in foil or painted to block passive light transmission.

### Double Tower Design

To accommodate environments with limited vertical clearance, a Double Tower version of the system was developed (Figure 3; Tables 3–4). This design allows two towers to be used within the same footprint as a single tower, maximizing use of both horizontal and vertical space. The lid was modified to support two towers, and specialized 3- and 4-cup modules were designed to ensure adequate light exposure for all plants. To maintain equal distribution between towers, the system uses a T-junction fitting at the pump outlet to split flow into two channels.

**Figure 3.**
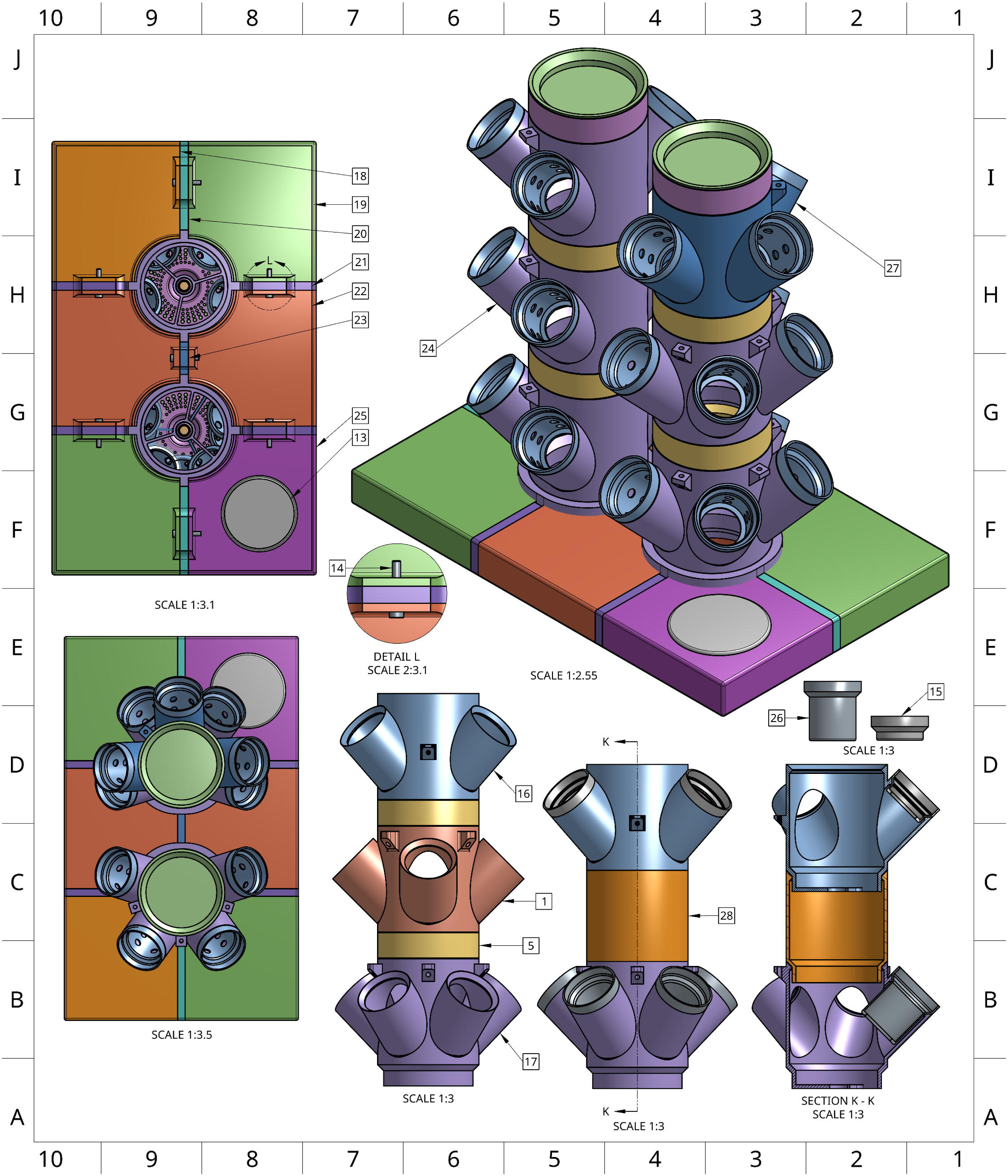
Multi-perspective rendering of the assembled Double Tower system and unique Double Tower components. An interactive 3D rendering of the Double Tower system (https://sketchfab.com/3d-models/two-tower-hydroponics-assembly-variant-models-d80f772011a44d1e928962e8bdacf65a) is available for close examination of parts in the context of a completed system.

**Table 3.** Tower modules for the double tower (DT) system.

| Part Number | File Name | Part Name | Count for one system |
| --- | --- | --- | --- |
| 24 | DT_4-Cup_Planting_Module.stl | 4-cup DT Planting Module | Variable |
| 2 | Tower_Lid.stl | Tower Lid | 2 |
| 3 | Tower_Distributor.stl | Tower Distributor | 2 |
| 4 | Stream_Breaker.stl | Stream Breaker | 2 |
| 5 | Tower_Short_Spacer.stl | Tower Short Spacer | Variable |
| 6 | Standard_Pot.stl | Standard Pot | 4 x 4-cup DT Planting Module |
| 7 | Reservoir_Tower_Adaptor.stl | Tower Adaptor | 2 |

**Table 4.** Lid modules for the double tower (DT) system.

| Part Number | File Name | Part Name | Count for one system |
| --- | --- | --- | --- |
| 13 | Access_Hole_Cap | Access Hole Cap | 1 |
| 18 | Reservoir_Lid_DT_Right.stl | Reservoir Lid DT Right | 1 |
| 19 | Reservoir_Lid_DT_Left.stl | Reservoir Lid DT Left | 2 |
| 20 | DT_Spacer_Long_Edge.stl | DT Spacer Long Edge | 4 |
| 21 | DT_Spacer_Short_Edge.stl | DT Spacer Short Edge | 2 |
| 22 | Reservoir_Lid_DT_Middle.stl | Reservoir Lid DT Middle | 2 |
| 23 | DT_Spacer_Middle_Edge.stl | DT Spacer Middle | 1 |
| 25 | Reservoir_Lid_DT_Right_Access_Hol.stl | Reservoir Lid DT Right Access Hole | 1 |
| 14 | Not applicable | M5 Screw and nut - 25 mm | 7 |

Each vinyl tubing line is extended by an additional ∼25 cm to accommodate the separate vertical stacks, allowing a single pump to serve both towers.

### Pre-Print and Pre-Assembly Recommendations

#### Print Specification

When printing parts for assembly, we recommend at least 20% infill to ensure that the lid and its components are structurally sound enough to support the tower modules and plants. Parts were fabricated using the Prusa MK3S+, that features a bedspace and headspace (minimum 250 × 210 × 210 mm). Using a printer with these dimensions, or larger, ensures that parts can be printed in single pieces and are not forced to bifurcate due to size constraints. While our 3D components were printed using PLA, other filament materials such as polyethylene terephthalate glycol (PETG) may offer provide superior stability and waterproofing for some applications.

#### Planting Module Compatibility

Planting modules are cross-compatible within their intended orientation: Single Tower or Double Tower components can be installed in any order or rotated in any direction. However, while modules designed for the Double Tower are compatible with the Single Tower setup, it is not recommended to use Single Tower modules in the Double Tower configuration. Single tower modules do not allow for sufficient space near each plant when installed in a double tower configuration.

#### Basic Sizing Overview

When considering location and space limitation for your assembled device, refer to Figure 4, which shows all major dimensions for a standard Single Tower setup (three planting/spacer modules stacked). These sizing dimensions remain the same for the Double Tower variant, as both versions share the same height, width, and length. An interactive 3D rendering of the Single Tower system (https://sketchfab.com/3d-models/one-tower-enviroment-696d529bdf3249bb8fe916f2aff49538) and Double Tower system (https://sketchfab.com/3d-models/two-tower-hydroponics-assembly-variant-models-d80f772011a44d1e928962e8bdacf65a) are available for close examination of parts in the context of a completed system.

**Figure 4.**
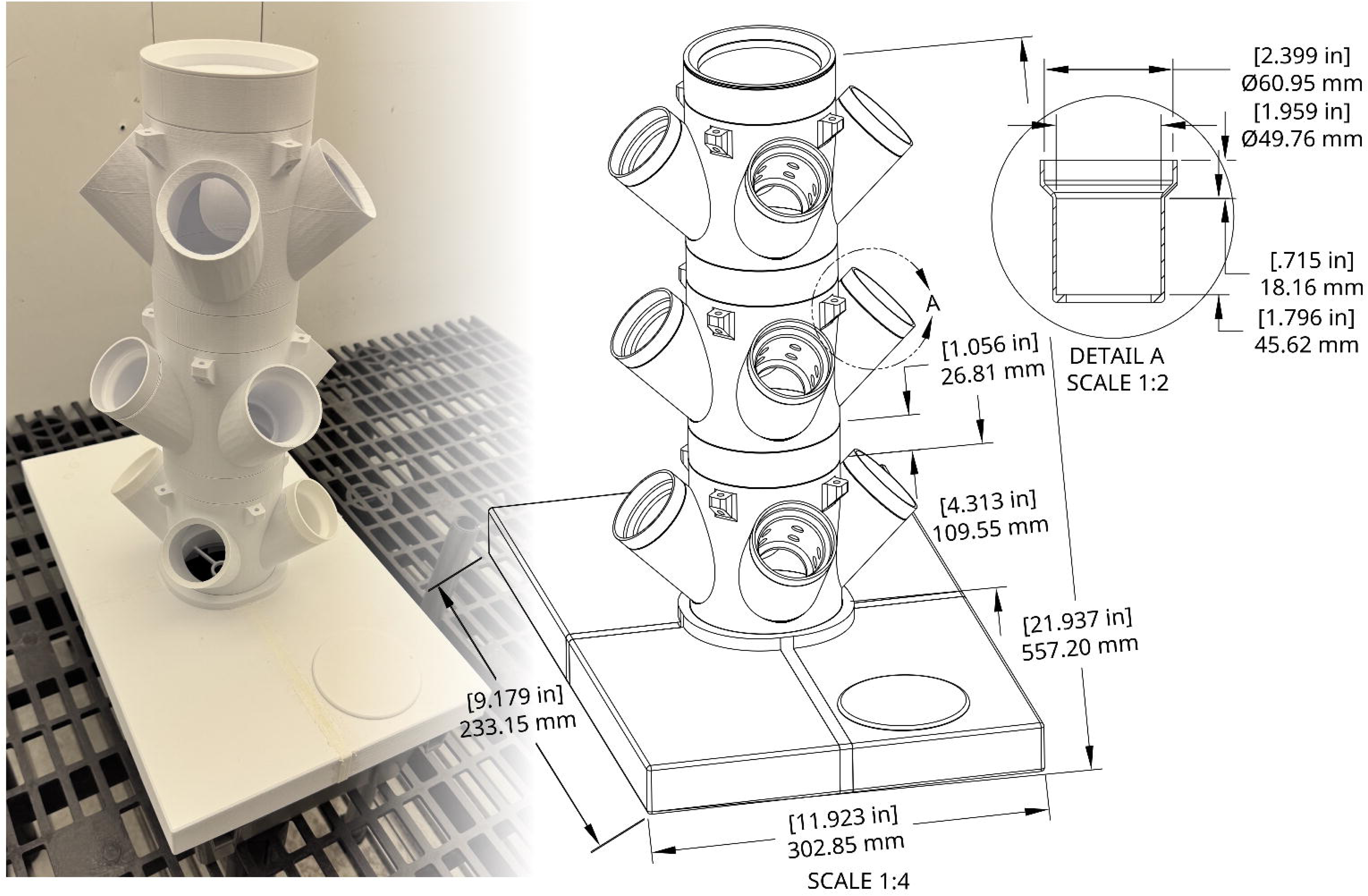
Photograph of an assembled Single Tower system with accompanying rendering showing dimensions of each component.

**Figure 5.**
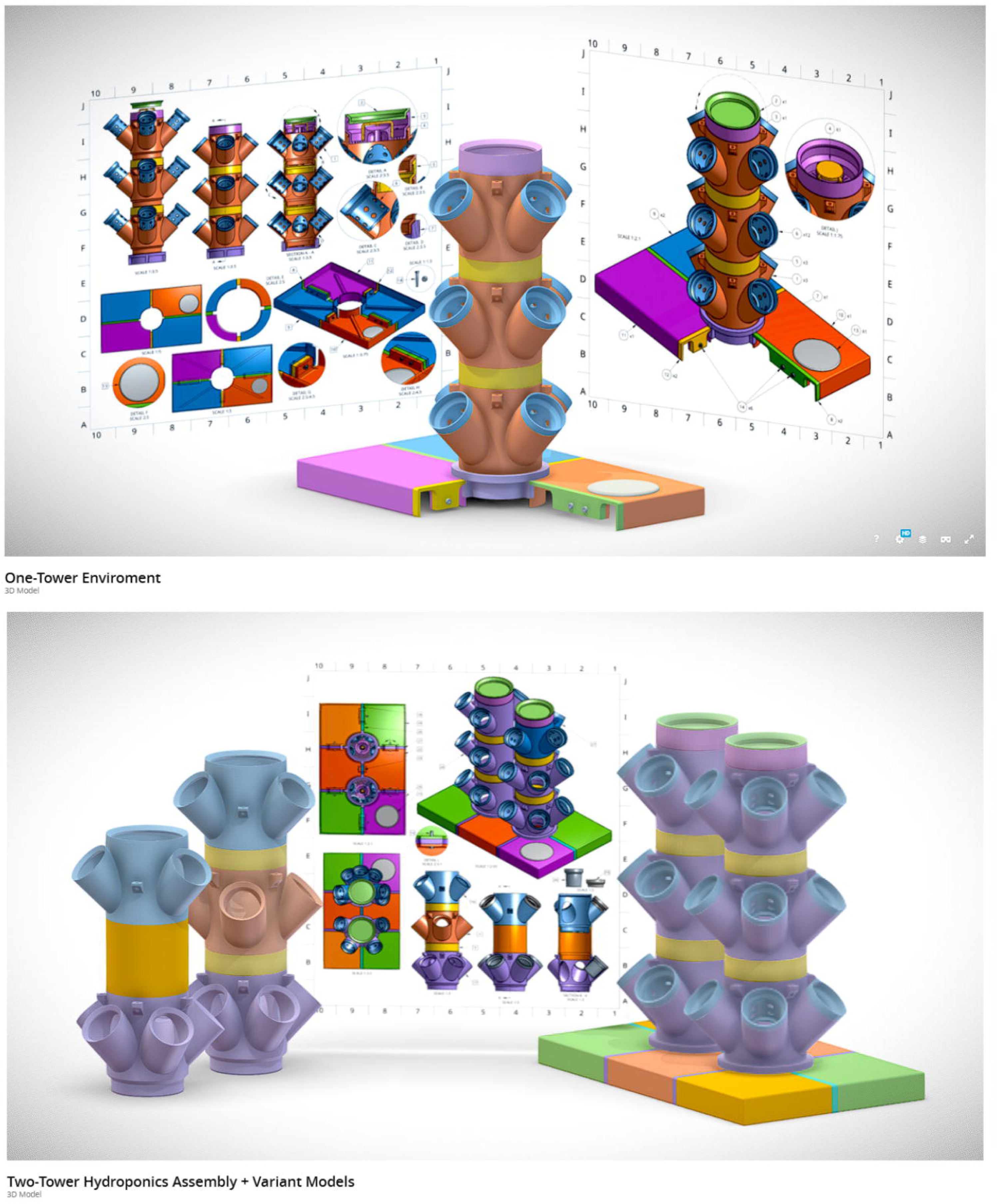
Preview display of the interactive render environments for the Single and Double Tower Assembly variations. An interactive 3D rendering of the Single Tower system (https://sketchfab.com/3d-models/one-tower-enviroment-696d529bdf3249bb8fe916f2aff49538). An interactive 3D rendering of the Double Tower system (https://sketchfab.com/3d-models/two-tower-hydroponics-assembly-variant-models-d80f772011a44d1e928962e8bdacf65a)

#### Core Device Assembly

*For details on device assembly please refer to the protocol document on Protocol.IO* and included in the Supporting Information. Variations for plant cups, planting modules, and spacers that have alternative designs and dimensions are listed in Table 5.

**Table 5.** Part variations for all systems.

| Part Number | File Name | Part Name | Count for one system |
| --- | --- | --- | --- |
| 15 | Short_Pot.stl | Short Pot | NA |
| 16 | ST_3-Cup_Planting_Module.stl | 3-cup ST Planting Module | NA |
| 17 | ST_5-Cup_Planting_Module.stl | 5-cup ST Planting Module | NA |
| 26 | True_Standard_Pot.stl | True Standard Pot | NA |
| 27 | DT_3-Cup_Planting_Module.stl | 3-cup DT Planting Module | NA |
| 28 | Tower_Tall_Spacer.stl | Tower Tall Spacer | NA |

### Operational Considerations

#### Solution Considerations

This hydroponics system is designed to function with 15L of solution in the reservoir. Lower volumes of solution may result in incomplete submersion of the hydroponics pump and would necessitate more frequent changes as plants matured. For our experiment we used Hoagland’s No. 1 nutrient solution [28]. However, any hydroponics solution is foreseeably compatible with this system. We recommend conducting preliminary experiments with the crops of interest to determine the optimal frequency by which the nutrient solution is replaced.

#### Pump Considerations

As referenced earlier in the paper, we used an ActiveAqua Submersible Pump 250 for our experiment. However, any submersible hydroponics pump will work provided it has sufficient power to carry nutrient solution to the top of the tower, and can be connected to the ½” inner-diameter tubing. Furthermore, we adjusted the flow rate of the pumps to approximately 200 gallons per hour. This flow rate is likely sufficient for most leafy greens, but may need to be optimized for other crops by the end user.

#### Planting Considerations

When planting, plants should be placed into 2” neoprene collars or a media, such as expanded clay pellets, to ensure they do not shift out of position. Our recommendation is to use a slant board system and transplant seedlings into neoprene collars and install into plant cups after cotyledon emergence as well described by Langenfeld and Bugbee [29]. A 3D printed slant board mount can be found here (https://3d.nih.gov/entries/3DPX-021396) [30].We found that direct seeding into neoprene collars without pre-germinating on slant boards resulted in inconsistent emergence and growth.

#### Case Study

We conducted a pilot study using this hydroponics system to grow four varieties of spinach in different salinity conditions. Our methods and results are described below.

## Methods

### System construction

Files for the single tower were downloaded from the NIH3D database (https://3d.nih.gov/entries/3DPX-021941), converted into gcode using PrusaSlicer, and printed on a Prusa MK3S+ 3D printer with PLA filament at 20% infill [25]. 3D printed objects were assembled into three individual single tower systems with three levels of 4-plant modules as described in our detailed protocol (dx.doi.org/10.17504/protocols.io.e6nvw4pw9lmk/v2; Supporting Information).

### Plant material, salinity treatments, and harvest conditions

Four cultivars of spinach (Sun Angel F1, Red Snapper F1, Auroch F1, and Flamingo Improved F1) were used in this study. Seeds were osmoprimed between two pieces of Whatman filter paper for three days in petri dishes containing 47.5:1 water:hydrogen peroxide solution.

Petri dishes were placed in a Conviron PGW36 growth chamber (Conviron; Winnipeg, Canada) maintained at 20 °C and 55 ± 10 % relative humidity, with 200 µmol/m^2^/s of photosynthetically active radiation for 12 hours over 10 days. After three days, seedlings were transferred onto felt wicks and installed into 2” neoprene collars and installed in plant cups. A detailed protocol for spinach germination can be found here: dx.doi.org/10.17504/protocols.io.5qpvodxebg4o/v2.

Plants were randomly assigned into one of three blocks, which correspond with their height within the hydroponics towers they were installed into. Block 1 plants were on the lowest tower module, block 2 plants were on the middle tower module, and block 3 plants were on the top tower module. Plants were grown in quarter strength Hoagland’s No. 1 solution (0.6 dS/m) days 0-7 and switched to half-strength (1.2 dS/m) days 8-14. Full strength Hoagland’s No. 1 (2.4 dS/m) was provided to all plants days 14-21 until the start of treatments. For the remainder of the experiment (days 22-35), plants were provided either full strength Hoagland’s No. 1 nutrient solution (control), control + 20/10 mM NaCl/CaCl_2_ to an electrical conductivity (EC) of 4.6 dS/m (low salinity), or control + 20/10 mM NaCl/CaCl_2_ to an EC of 6.4 dS/m (mild salinity).

The pH of all nutrient solutions was maintained between 5.5 and 6.0 using 0.2 M H_2_SO_4_ and 0.1 M KOH to lower or raise pH, respectively. After 35 days when plants had 6-8 mature leaves, tissue was harvested, weighed, and immediately frozen at -80 °C before processing. Dry weight was determined by lyophilizing pre-weighed frozen tissue over five days and subtracting the calculated moisture content from the original sample mass.

### Statistical analysis and data visualization

Data were analyzed as a randomized complete block design with three treatments, four varieties, and three blocks. Analysis of variance was conducted in R statistical software followed by Tukey’s honestly significant difference test using the ‘car’ package [31,32]. Data were visualized using ggplot2 [33].

## Expected Results

### Fresh weight yield

There were no significant differences in mean fresh weight across any of the treatments when analyzing all varieties together (Figure 6; Table 6). Of the three treatments, mild salinity had the highest mean fresh weight at 26.23 grams, while the control treatment had the lowest at 24.09 grams. There was no significant treatment effect on fresh weight (p = 0.77), nor treatment-variety interaction (p = 0.95). However, the effect of variety on fresh weight was highly significant (p < 0.001), indicating that variety played a stronger role than salinity treatment on fresh weight. Auroch F1 had the highest mean fresh weight at 37.90 grams, while Red Snapper F1 had the lowest mean fresh weight at 7.95 grams. Flamingo F1 Improved and Sunangel F1 had mean fresh weight yields of 29.44 and 24.92 grams, respectively. Fresh weight yields are summarized in Table 1. Within each variety, there were no significant differences noted between treatment groups with the exception of Flamingo Improved F1, which had a significant difference between the Low and Mild salinity conditions (Table 6; Figure 6). However, there was statistical significance for differences in fresh weight yield between varieties. Red Snapper F1 was significantly different from Auroch F1 in the Control treatment, while in the Low salinity treatment it was significantly different from both Auroch F1 and Flamingo Improved F1. In the mild treatment, Sunangel F1 and Flamingo Improved F1 were significantly different.

**Figure 6.**
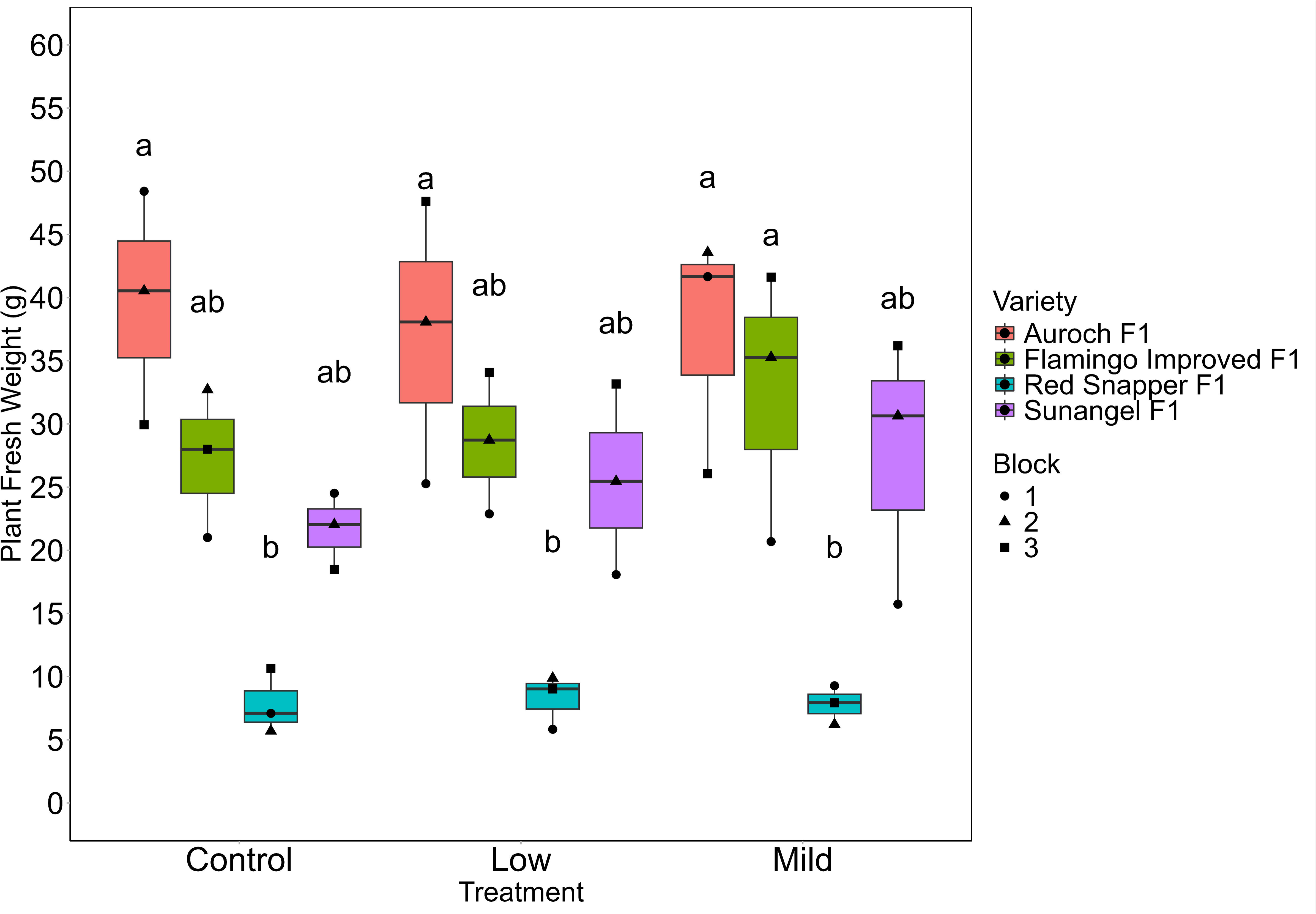
Fresh weight yield per plant by treatment and variety. Group letters from Tukey’s post- hoc test for significant differences (11=0.05) are displayed above each group.

**Table 6.**
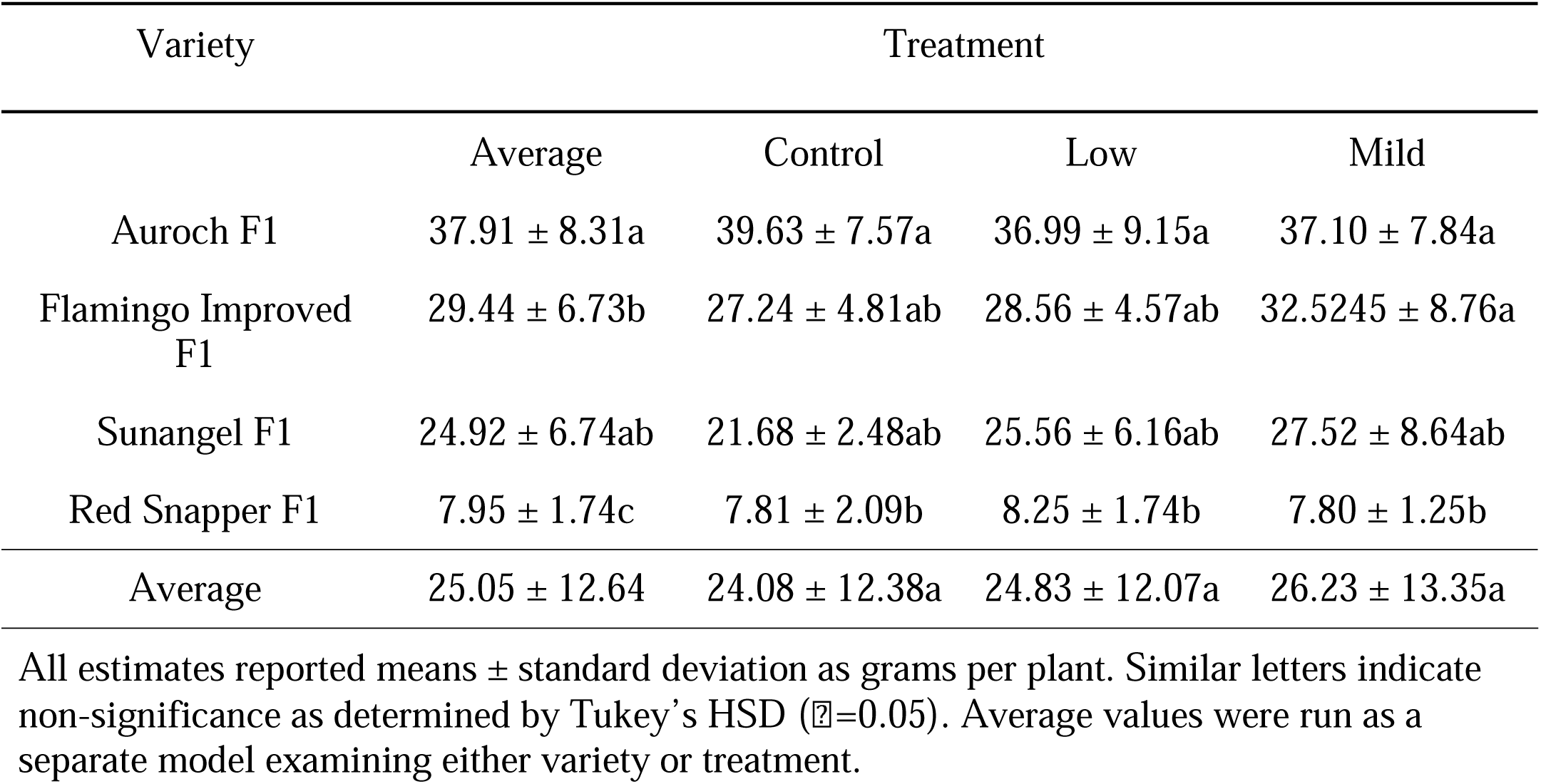
Mean fresh weight per plant (g) by treatment and variety.

| Variety | Treatment |  |  |  |
| --- | --- | --- | --- | --- |
|  | Average | Control | Low | Mild |
| Auroch F1 | 37.91 ± 8.31a | 39.63 ± 7.57a | 36.99 ± 9.15a | 37.10 ± 7.84a |
| Flamingo Improved F1 | 29.44 ± 6.73b | 27.24 ± 4.81ab | 28.56 ± 4.57ab | 32.5245 ± 8.76a |
| Sunangel F1 | 24.92 ± 6.74ab | 21.68 ± 2.48ab | 25.56 ± 6.16ab | 27.52 ± 8.64ab |
| Red Snapper F1 | 7.95 ± 1.74c | 7.81 ± 2.09b | 8.25 ± 1.74b | 7.80 ± 1.25b |
| Average | 25.05 ± 12.64 | 24.08 ± 12.38a | 24.83 ± 12.07a | 26.23 ± 13.35a |

All estimates reported means ± standard deviation as grams per plant. Similar letters indicate non-significance as determined by Tukey's HSD ( $\alpha=0.05$ ). Average values were run as a separate model examining either variety or treatment.

### Dry Weight Data

There were no significant differences in dry weight percentage per plant between any of the varieties or treatments (Table 7; Figure 7). Dry weight percentages varied based on variety, with Red Snapper F1 having a slightly higher dry weight percentage per plant when compared to Auroch F1, Flamingo Improved F1, and Sunangel F1. Dry weight values for each variety and treatment are summarized in Table 7.

**Figure 7.**
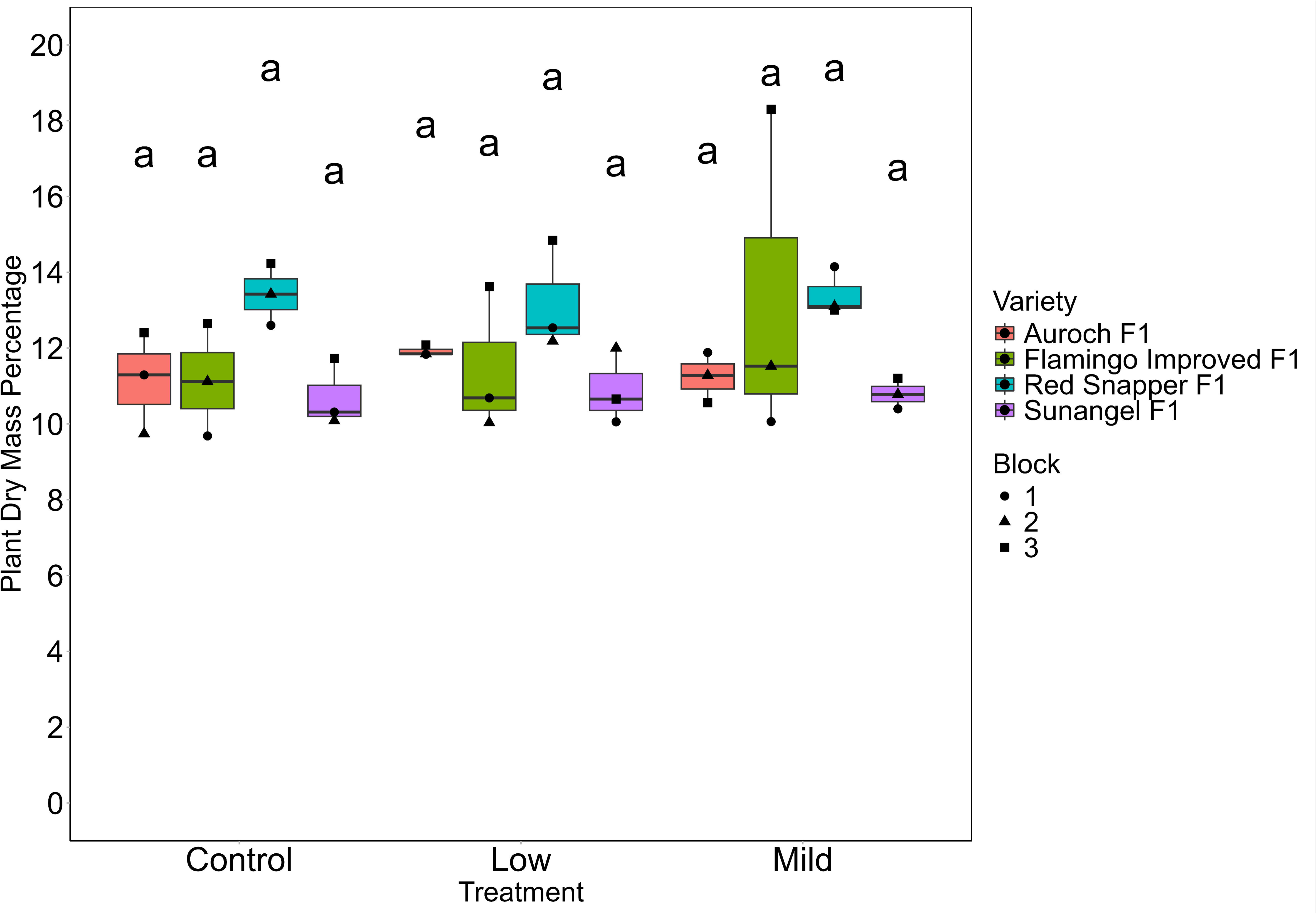
Dry mass percentage per plant by treatment and variety. Group letters from Tukey’s post-hoc test for significant differences (11=0.05) are displayed above each group.

**Table 7.** Mean dry weight percentage per plant by treatment and variety.

| Variety | Treatment |  |  |  |
| --- | --- | --- | --- | --- |
|  | Average | Control | Low | Mild |
| Auroch F1 | 11.12 ± 0.79ab | 11.15 ± 1.34a | 11.92 ± 0.14a | 11.24 ± 0.66a |
| Flamingo Improved F1 | 11.62 ± 2.55ab | 11.15 ± 1.48a | 11.45 ± 1.91a | 13.29 ± 4.40a |
| Sunangel F1 | 10.80 ± 0.66b | 10.71 ± 0.89a | 10.90 ± 1.00a | 10.79 ± 0.40a |
| Red Snapper F1 | 13.34 ± 0.84a | 13.42 ± 0.82a | 13.19 ± 1.44a | 13.42 ± 0.63a |
| Average | 11.72 ± 1.72 | 11.61 ± 1.43a | 11.71 ± 1.36a | 11.85 ± 2.19a |

All estimates reported means ± standard deviation. Similar letters indicate non-significance as determined by Tukey's HSD ( $\alpha=0.05$ ). Average values were run as a separate model examining either variety or treatment.

### Qualitative observations

Plants in the lower levels of the towers appeared to be noticeably smaller than their counterparts in the middle and upper levels of the towers. We hypothesized that this was due to shading effects from the plants above and distance from the light canopy, as the light source for this study was an overhead point source of lighting. We observed that the average photosynthetic flux density (PPFD) at the lowest, middle, and highest tower module was 355, 371, and 457 µmol/m^2^/s. Given a 12-hour photoperiod, this equates to a daily light integral (DLI) of 15.34, 16.04, and 19.74. The DLI of plants in block 3 (the top of our towers) was substantially higher due to being physically closer to the light fixture and not having shading from plants above.

Although there were no significant differences in plant fresh weight or dry weight percentage by block, block 3 plants had the highest fresh weights in 6 of the 12 unique combinations (Figure 6) and dry weight percentage and had the highest dry weights in 9 of the 12 unique salinity-variety combinations (Figure 7). This finding suggests that shading and distance from the light canopy could be a concern for researchers operating in growth chambers and we suggest alternative lighting strategies such as laterally placed fixtures or using natural sunlight for more uniform growth Additionally, at the end of the experiment the nutrient solution flow rate was noted to be lower for the control tower when compared to the low and mild salinity towers. During disassembly at harvest, significant root growth was observed in the filter and intakes of the submersible pumps in all bins. For longer duration experiments (e.g. > five weeks) we strongly recommend developing either a preventative or maintenance strategy to mitigate root ingress into nutrient solution pumps.

**Supporting information** S1: Step-by-step protocol, also available on protocols.io: dx.doi.org/10.17504/protocols.io.e6nvw4pw9lmk/v2. S2: Raw data used for figures presented in this study.

## Supporting information

Detailed protocol for hydroponics system

## Acknowledgements

We thank Elaine Le and Alvin Tak for assistance with plant maintenance and harvest.

## Authors’ Contributions

Conceptualization: Michael Dzakovich, Suraj Chandramouli, and Ethan Shaw Investigation: Suraj Chandramouli Writing – original draft: Suraj Chandramouli, Ethan Shaw, and Michael Dzakovich Writing – review & editing: Ethan Shaw, Michael Dzakovich, and Suraj Chandramouli

## Author disclaimer

The findings and conclusions in this publication are those of the authors and should not be construed to represent any official USDA or U.S. Government determination or policy. Mention of trade names or commercial products in this publication is solely for the purpose of providing specific information and does not imply recommendation or endorsement by the U.S. Department of Agriculture. The USDA is an equal opportunity provider and employer.

## Data availability

All data associated with this manuscript are presented within and included in the supporting information.

## Associated Content

A step-by-step protocol is available on protocols.io: dx.doi.org/10.17504/protocols.io.e6nvw4pw9lmk/v2.

