## Supplementary material for "Layers to Leaves: A Suite of Modular 3D Printed Hydroponics Components for Research and Education": Detailed protocol for hydroponics system

Jul 04, 2025 Version 2

### Dzakovich Lab 3D Printed Hydroponics System Setup V.2

DOI
dx.doi.org/10.17504/protocols.io.e6nvw4pw9lmk/v2

Ethan Shaw[1], Suraj Chandramouli[2], Michael Dzakovich[3]
[1]University of Houston; [2]Rice University; [3]USDA-ARS Children's Nutrition Research Center

Dzakovich Lab

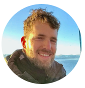

Michael Dzakovich
USDA-ARS Children's Nutrition Research Center

OPEN ACCESS

**DOI: dx.doi.org/10.17504/protocols.io.e6nvw4pw9lmk/v2**

**Protocol Citation:** Ethan Shaw, Suraj Chandramouli, Michael Dzakovich 2025. Dzakovich Lab 3D Printed Hydroponics System Setup. **protocols.io https://dx.doi.org/10.17504/protocols.io.e6nvw4pw9lmk/v2**Version created by **Michael Dzakovich**

**License:** This is an open access protocol distributed under the terms of the **Creative Commons Attribution License**, which permits unrestricted use, distribution, and reproduction in any medium, provided the original author and source are credited

**Protocol status:** Working
**We use this protocol and it's working**

**Created:** July 03, 2025

**Last Modified:** July 04, 2025

**Protocol Integer ID:** 221774

**Keywords:** 3D printing, Hydroponics, Research, Education, Plant biology, STEM

**Funders Acknowledgements:**
USDA-ARS CRIS Funds
Grant ID: 3092-10700-066-001S

#### Abstract

This protocol provides step by step instructions needed to assemble the Dzakovich Lab's Single Tower and Double Tower 3D printed hydroponics systems.

#### Materials

3D printer (FDM, PLA and/or PETG-compatible) — for fabricating all printed components from .STL files
-Single Tower: https://3d.nih.gov/entries/3DPX-021941
- Double Tower: https://3d.nih.gov/entries/3DPX-021942
M5 screws (25 mm length) — 6 required for Single-Tower, 8 for Double-Tower
M5 nuts — 6 required for Single-Tower, 8 for Double-Tower
M5 Allen wrench — for tightening screw heads during lid assembly
Adjustable wrench — for securing nuts during tightening
Silicon sealant — for sealing seams between lid and spacer modules
Silicon sealant extruder/tube — for applying sealant
Aluminum foil or dark opaque paint — optional, for light proofing lid
Box cutter or similar cutting tool — for trimming vinyl tubing and modifying bin
Flexible vinyl tubing (½″ inner diameter) — used for nutrient solution flow; exact length depends on tower height
Hydroponic pump — user’s choice, placed at bottom of bin
250-500-gram weight — (optional) used to secure Tower Lid module after assembly
Storage bin (e.g., ULine S-20588GR) — used as the base reservoir
Rotary tool with cutoff disc — used to remove bin lid hinges and rim tabs
Foam weather stripping (¾″ wide × 7/16″ thick) — for lining the bin rim
Soft cloth or sponge — for cleaning interior bin surfaces
PVC T-joint (½″ inner diameter) — required only for Double-Tower tubing setup

#### Bin Preparation (ULine S-20588GR)

1 Remove the original bin lid by clipping the hinges with wire cutters and discarding or repurposing the lid pieces.

2 Remove plastic tabs on the rim of each bin using a rotary tool with a cutoff cutting disc.

3 Cut a notch approximately 2 cm by 4 cm out of the rim along one side of the bin (confirm notch alignment with the figure in step 4) for the pump power cord.

4 Line the rim of each bin with 3/4″ wide x 7/16″ thick foam weather stripping.

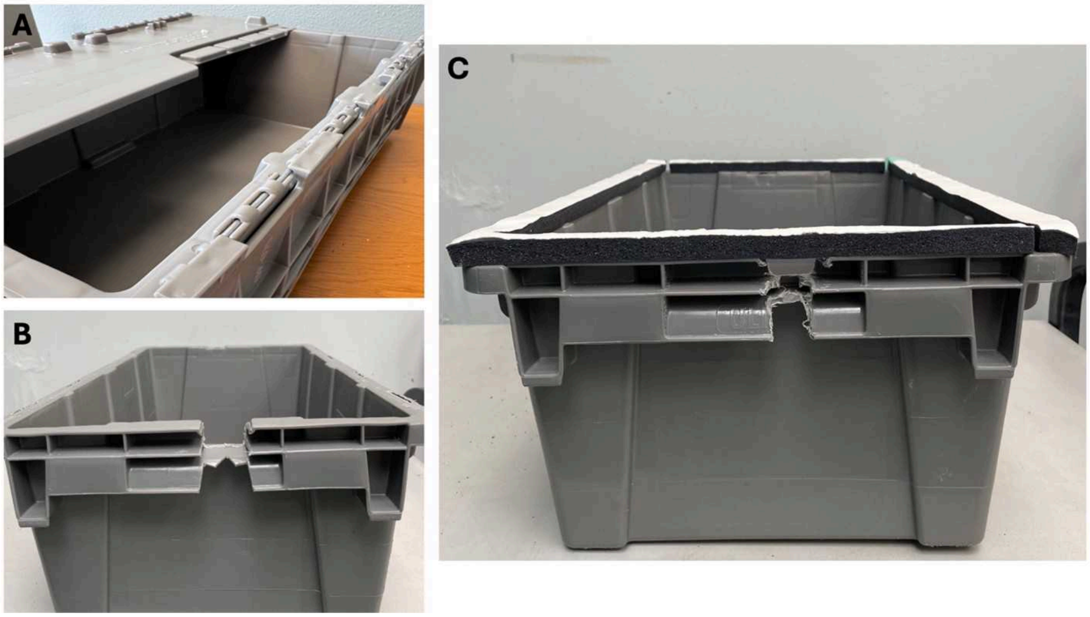


5 Wipe the inner surfaces of the bin to remove any stray plastic that could interfere with nutrient solution pumps with a soft cloth or sponge to avoid scratching the plastic surface.

#### Printing Setup and Specifications Directory

6 Download all files from the 3D-parts (.STL) directory.

7 Import files into a 3D-Slicer of choice and set infill to 20%.

8 For Single-Tower printing instructions follow steps 10-11, and for Double-Tower printing instructions follow steps 12-13.

#### Printing Instructions (Single-Tower)

9 Slice and print the following for the Single-Tower Lid: x2 Reservoir Spacer Long-Edge (Reservoir_Spacer_Long Edge.stl; Part 8), x2 Reservoir Spacer Short-Edge (Reservoir_Spacer_Short_Edge.stl; Part 12), x2 Reservoir Lid Left (Reservoir Lid_Left.stl; Part 9), x1 Reservoir Lid Right Access Hole (Reservoir_Lid Right _Access_Hole.stl; Part 10), x1 Reservoir Lid Right (Reservoir Lid Right.stl; Part 11), and x1 Access Hole Cap (Access Hole Cap; Part 13).

10 Slice and print the following for a standard 3-module-stacked Planting Tower: x3 4-cup ST Planting Module (4-Cup_Planting_Module.stl; Part 1), x2 Tower Short Spacer (Tower_Short_Spacer.stl; Part 5), x12 Standard Pot (Standard_Pot.stl; Part 6), x1 Stream Breaker (Stream_Breaker.stl; Part 4), x1 Tower Distributor (Tower_Distributor.stl; Part 3), x1 Tower Lid (Tower_Lid.stl; Part 2), and x1 Tower Adaptor (Reservoir_Tower_Adaptor.stl; Part 7).

#### Printing Instructions (Double-Tower)

11 Slice and print the following for the Double-Tower Lid: x1 Reservoir Lid DT Right (Reservoir_Lid_DT_Right.stl; Part 18), x2 Reservoir Lid DT Left (Reservoir_Lid_DT_Left.stl; Part 19), x2 DT Spacer Long-Edge (DT_Spacer_Long_Edge.stl; Part 20), x4 DT Spacer Short-Edge (DT_Spacer_Short_Edge.stl; Part 21), x2 Reservoir Lid DT Middle (Reservoir_Lid_DT_Middle.stl; Part 22), x2 DT Spacer Middle (DT_Spacer_Middle_Edge.stl; Part 23), x1 Reservoir Lid DT Right Access Hole (Reservoir_Lid_DT_Right_Access_Hole.stl; Part 25).

12 Slice and print the following for a standard 3-module-stacked Planting Tower: x6 4-cup DT Planting Module (DT_4-Cup_Planting_Module.stl; Part 24), x4 Tower Short Spacer (Tower_Short_Spacer.stl; Part 5), x24 Standard Pot (Standard_Pot.stl; Part 6), x2 Stream Breaker (Stream_Breaker.stl; Part 4), x2 Tower Distributor (Tower_Distributor.stl; Part 3), x2 Tower Lid (Tower_Lid.stl; Part 2), and x2 Tower Adaptor (Reservoir_Tower_Adaptor.stl; Part 7).

#### Assembly Specifications Directory

13 For Lid assembly instructions involving the Single-Tower design refer to steps 16-44, for Double-Tower design refer to steps 68-90.

14 For Tower assembly and tubing routing instructions involving the Single Tower design refer to steps 45-67, for Double-Tower design refer to steps 89-106.

#### Lid Assembly Instructions (Single-Tower)

15 Gather the printed components from step 10 and the Tower Adaptor (Part 7) from step 11.

16 Gather 6 × M5 screws (25-mm long) and 6 × M5 nuts.

17 Gather an M5 Allen wrench, Silicon Sealant Tube/Extruder, and either a sheet of aluminum foil or dark paint.

18 Place all lid modules upside down on a flat surface.

19 Arrange the lid modules in a 2×2 configuration: Top-left corner: Reservoir Lid Left (Part 9), Top-right corner: Reservoir Lid Right (Part 11), Bottom-left corner: Reservoir Lid Right Access Hole (Part 10), Bottom-right corner: Reservoir Lid Left (Part 9).

20 Insert one Long Edge Spacer (Part 8) between the top-left and top-right lid modules (along the top edge, long axis).

21 Insert the second Long Edge Spacer (Part 8) between the bottom-left and bottom-right lid modules (along the bottom edge, long axis).

22 Insert one Short Edge Spacer (Part 12) between the top-left and bottom-left lid modules (along the left edge, short axis).

23 Insert the second Short Edge Spacer (Part 12) between the top-right and bottom-right lid modules (along the right edge, short axis).

24 Align all bolting tabs on the underside of the lid and spacer modules where they meet.

25 Firmly press together each pair of adjacent lid and spacer modules to ensure tight contact.

26 Insert an M5 screw through each of the 5 aligned bolting tab holes (where lid modules meet spacers).

27 Thread an M5 nut onto the exposed end of each screw from below the assembly.

28 Use an M5 Allen wrench to hold each screw head in place.

29 Use an adjustable wrench to tighten each nut to hand-tightness (do not over-tighten).

30 Locate the Tower Adaptor module (Part 7).

31 Insert the Tower Adaptor (Part 7) into the central “cross” opening formed by the four lid modules.

32 Ensure the Tower Adaptor is seated evenly and flush with the surrounding lid.

33 Confirm you have completed the module correctly by referring to the figure below.

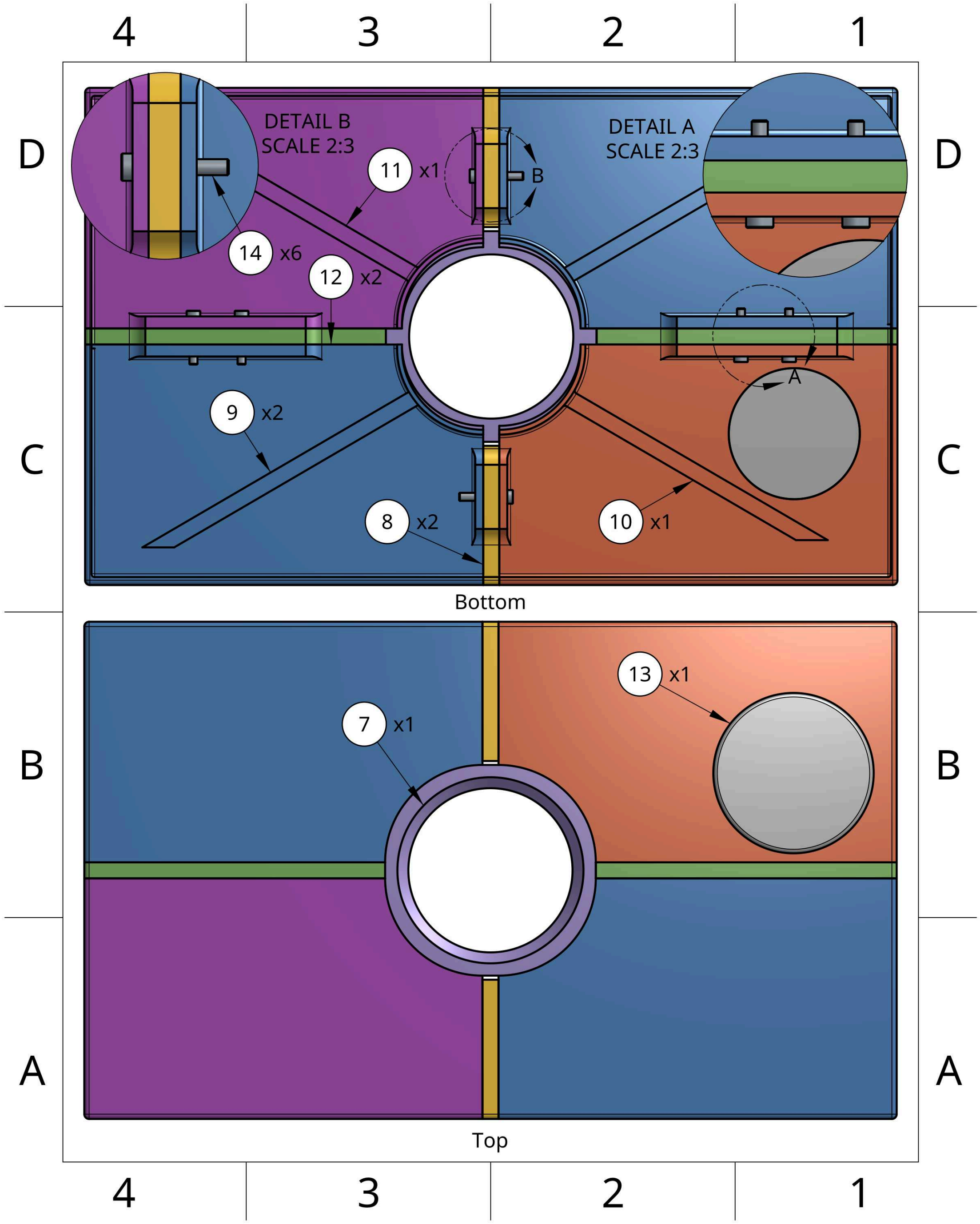


34 Prepare a silicon sealant bead approximately 1-mm thick.

35 Apply silicon sealant along all interior joints between the lid modules and spacer modules.

36 Apply silicon sealant along all exterior joints on the outside faces where modules meet.

37 Apply sealant around the outer radial seam where the Tower Adaptor (Part 7) interfaces with the lid surface.

38 Allow sealant to cure per manufacturer's recommendations before further handling.

39 Lightproof the lid by one of the following methods: Cover the top surface of the lid with aluminum foil; Paint the lid using dark-colored, opaque paint.

39.1 Alternatively, print lid components using a dark filament.

40 Locate the Access Hole Cap module (Part 13).

41 Place the Access Hole Cap over the access hole (on the Reservoir Lid Right Access Hole module (Part 10)).

42 Ensure the cap fits snugly to prevent solution evaporation and block ambient light.

#### Tower Assembly and Tubing Instructions (Single-Tower)

43 Gather the printed components from step 11.

44 Gather a box cutter (or similar cutting tool), a spool of ½" inner diameter flexible vinyl tubing, a ~250-500-gram weight, and the user's choice of pump.

45 Place the base Planting Module (Part 1) into the Tower Adaptor (Part 7) on top of the lid.

46 Attach a Tower Spacer Module (Part 5) on top of the base Planting Module (Part 1).

46.1 Spacer modules are optional and available in multiple sizes to suit the user's needs.

47 Stack the next Planting Module (Part 1) on top of the spacer (Part 5).

48 Repeat steps 48-49 an additional time.

49 Once stacking is complete, place the Tower Distributor Module (Part 3) on top of the final Planting Module (Part 1).

50 Insert Planting Cups (Part 6) into the cavities along the Planting Modules (Part 1).

51 Confirm the setup is correct by referring to the figure below.

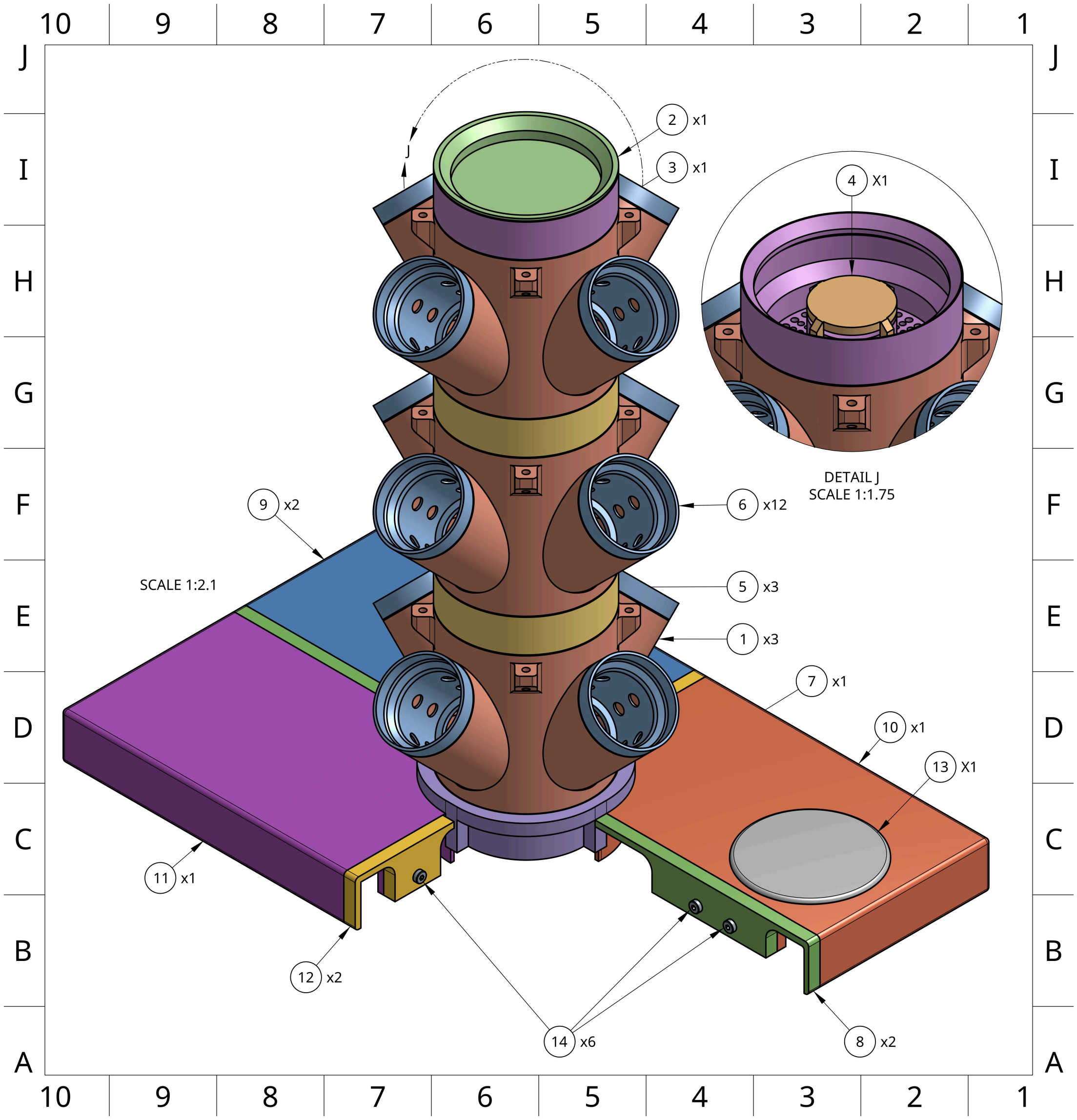


52 Measure the total height of the tower stack from the top of the Tower Distributor (Part 3) down through the modules to the bin floor.

53 Add 15-cm to the measured height to account for internal routing and flexibility.

54 Cut the ½ inch vinyl tubing to this length.

55 Connect the top end of the tubing to the nipple fitting located on the underside of the Tower Distributor Module (Part 3).

56 Thread the tubing downward through the central brackets of each module — passing through each Tower Spacer (Part 5) and Planting Module (Part 1) until it exits from the bottom of the tower.

57 Place the hydroponics pump into the bin such that the output nozzle is centered on the bin floor.

58 Route the power cable of the pump through the notch cut in the bin rim.

59 Lower the entire lid + tower assembly into position on the bin, aligning it securely with the bin walls.

60 Connect the bottom end of the vinyl tubing to the output nozzle of the pump.

61 Check for proper slack and avoid kinking: If the tubing is too long, disconnect it from the pump, trim the excess, and reconnect; Ensure the tubing lies without tension and allows the pump to sit flat.

62 Place the Stream Breaker Module (Part 4) on top of the Tower Distributor Module (Part 3).

63 Place the Tower Lid Module (Part 2) on top of the Stream Breaker (Part 4).

64 Add a weight (~250-500 grams) on top of the Tower Lid (Part 2) to secure the upper modules against displacement from water pressure.

64.1 Alternatively, seal lid with thin layer of silicone. A razor blade or other sharp object can cut through the silicone seal for system deconstruction and maintenance.

#### Lid Assembly Instructions (Double-Tower)

65 Gather the printed components from step 12 and the two Tower Adapter modules from step 13.

66 Gather 8 × M5 screws (25-mm long) and 8 × M5 nuts.

67 Place all lid modules upside down on a flat surface with bolting tabs facing up.

68 Arrange the 6 lid panels in a 2×3 configuration: Top Row: Reservoir Lid DT Left (Part 19) → Reservoir Lid DT Middle (Part 22) → Reservoir Lid DT Right (Part 18); Bottom Row: Reservoir Lid DT Left (Part 19) → Reservoir Lid DT Middle (Part 22) → Reservoir Lid DT Right Access Hole (Part 25).

69 Insert one Short Edge Spacer (Part 20) between the top and bottom DT Left modules (vertical joint).

70 Insert the second Short Edge Spacer (Part 20) between the top and bottom DT Middle modules (vertical joint).

71 Insert one Long Edge Spacer (Part 21) between the top-left and top-middle panels (horizontal joint).

72 Insert a second Long Edge Spacer (Part 21) between the top-middle and top-right panels (horizontal joint).

73 Insert a third Long Edge Spacer (Part 21) between the bottom-left and bottom-middle panels (horizontal joint).

74 Insert the fourth Long Edge Spacer (Part 21) between the bottom-middle and bottom-right panels (horizontal joint).

75 Insert the DT Spacer Middle (Part 23) between the two DT Middle panels (between the towers, center seam).

76 Align all tab holes on lid modules and spacer blocks.

77 Insert 8 × M5 Screws (Part 14) through the bolting tabs at each spacer joint.

78 Firmly press adjacent components together before threading nuts.

79 Thread M5 nuts onto the underside of each screw.

80 Tighten each screw to hand-tight using an M5 Allen wrench and adjustable wrench (do not over-tighten).

81 Locate the two Tower Adaptor modules (Part 7).

82 Insert one of the Tower Adaptors (Part 7) into the left-most “cross” opening, and insert another into the adjacent opening to the right.

83 Ensure the Tower Adaptor is seated evenly and flush with the surrounding lid.

84 Confirm you have completed the module correctly by referring to the figure below.

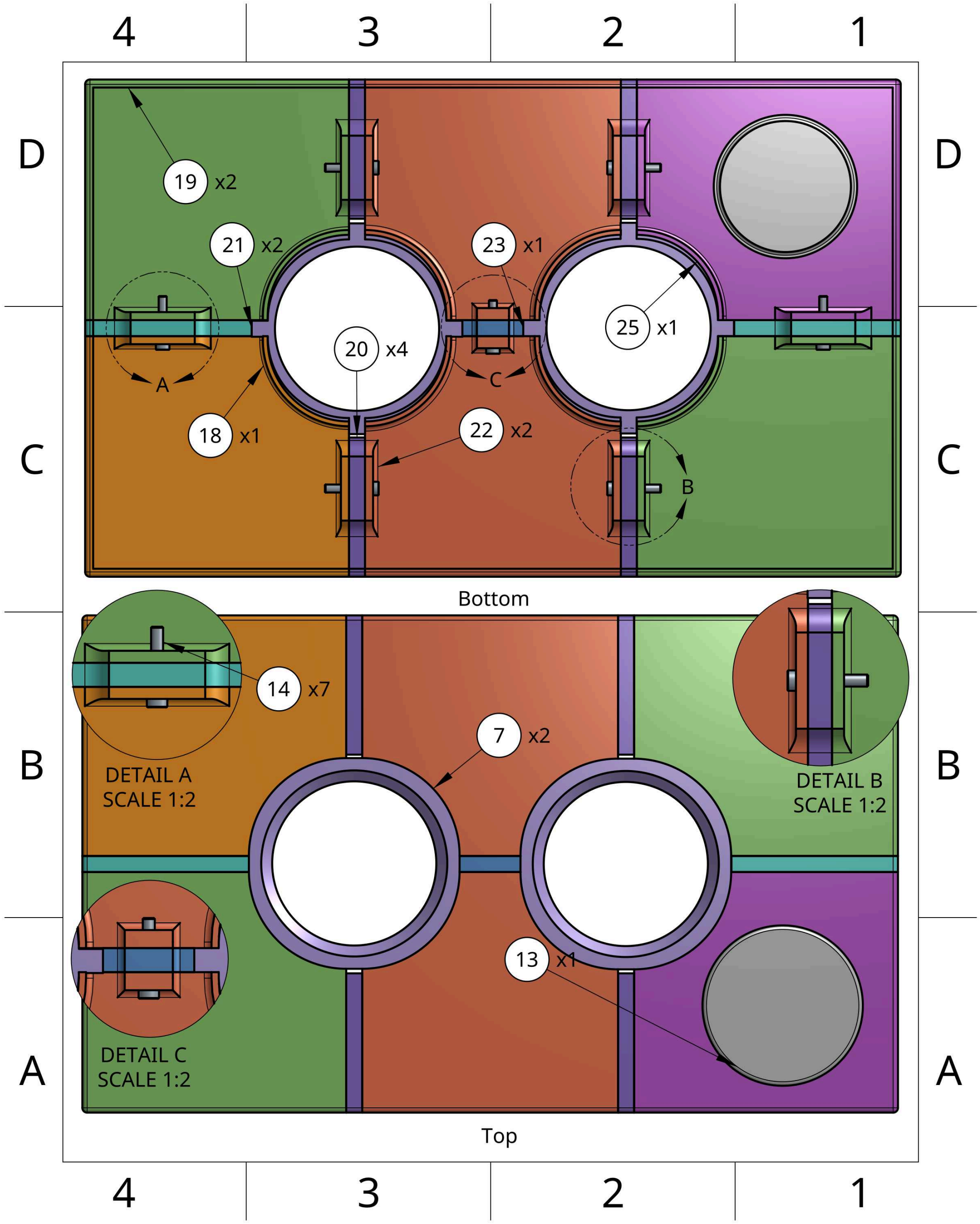


85 Complete the operations using silicon sealant from steps 36-40.

85.1 Optionally light proof with aluminum foil or print components with dark filament.

86 Place the Access Hole Cap (Part 13) into the access hole on the bottom-right lid module (Part 25).

#### Tower Assembly and Tubing Instructions (Double-Tower)

87 Gather the printed components from step 13.

88 Gather a box cutter (or similar cutting tool), a spool of ½″ inner diameter flexible vinyl tubing, a ~250-500-gram weight, the user’s choice of pump, and a ½″ inner diameter PVC T-Joint.

89 Complete the tower assembly operations dictated throughout steps 47-52 for both Tower Modules.

90 Confirm the setup is correct by referring to the figure below.

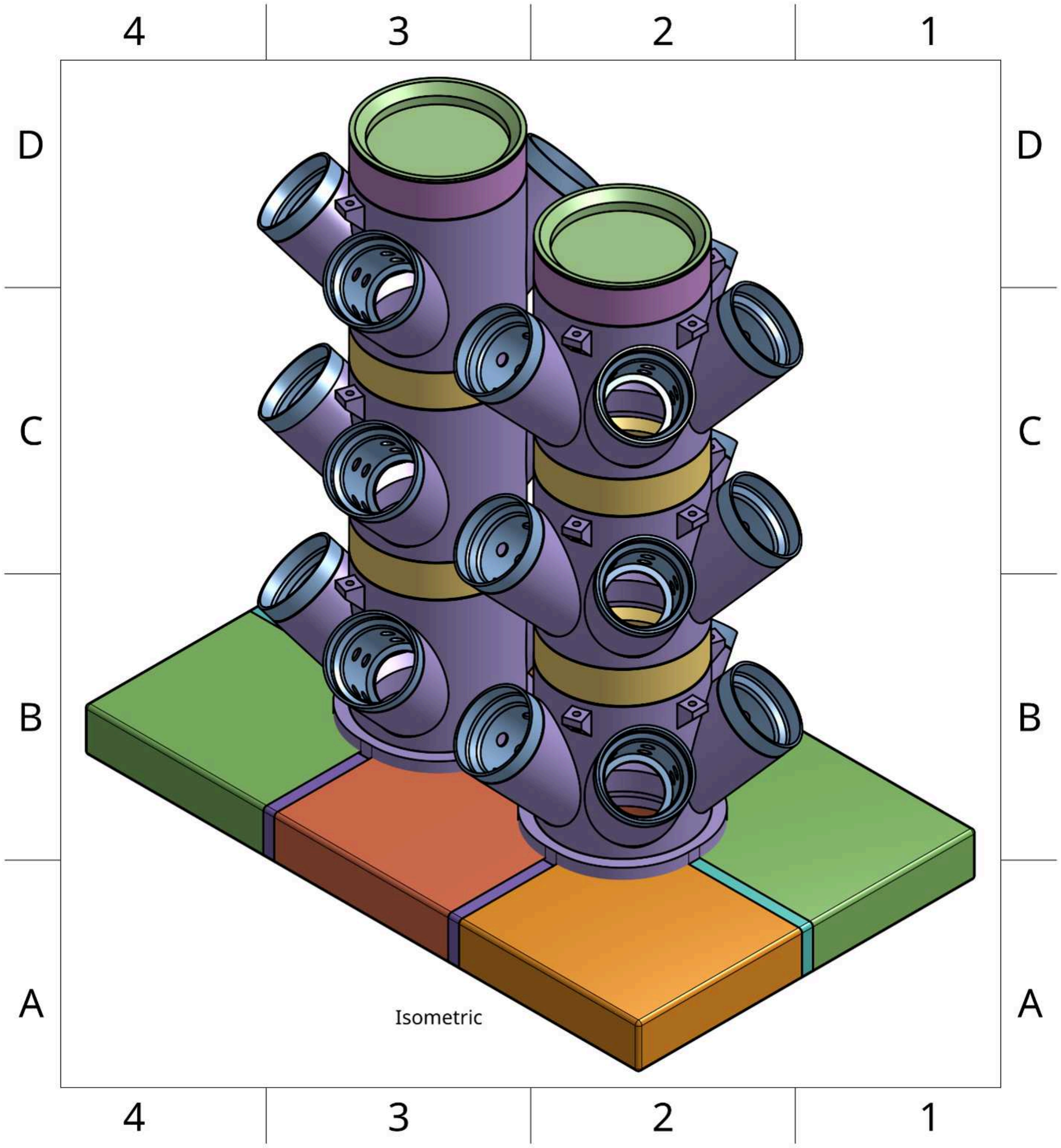


91 Measure the total height of the tower stack from the top of the Tower Distributor (Part 3) down through the modules to the bin floor.

92 Cut 2 equal lengths of ½ inch inner diameter vinyl tubing — each should be total tower height with an additional 25 cm.

93 Connect the top end of each tubing piece to the nipple fitting underneath each Tower Distributor Module (Part 3).

94 Thread each tubing piece downward through the tubing brackets inside its respective tower.

95 Let each tubing line exit cleanly at the bottom end of its tower.

96 Attach both tubing lines to the ½″ t-fitting, ensuring equal length from both towers to the junction.

97 Cut a short third piece of tubing from the same material — long enough to reach from the t-fitting to the pump output nozzle.

98 Connect this third piece to the bottom outlet of the t-fitting.

99 Complete the pump assembly operations dictated throughout steps 60-62.

100 Connect the short tubing line from the t-fitting to the output nozzle of the pump.

101 Check that tubing lengths allow the pump to sit flat at the bottom without kinking.

102 If tubing is too long, trim the excess and reconnect all ends.

103 Complete final operations dictated throughout steps 65-67.
